# Effect of structural stability on endolysosomal degradation and T-cell reactivity of major shrimp allergen tropomyosin

**DOI:** 10.1101/2020.02.17.919845

**Authors:** Sandip D. Kamath, Sandra Scheiblhofer, Christopher M. Johnson, Yoan Machado, Thomas McLean, Aya C Taki, Paul A. Ramsland, Swati Iyer, Isabella Joubert, Heidi Hofer, Michael Wallner, Josef Thalhamer, Jennifer Rolland, Robyn O’Hehir, Peter Briza, Fatima Ferreira, Richard Weiss, Andreas L. Lopata

## Abstract

**Background:** Tropomyosins are highly conserved proteins, an attribute that forms the molecular basis for their IgE antibody cross-reactivity. Despite structural similarities, their allergenicity varies greatly between ingested and inhaled invertebrate sources. In this study, we investigated the relationship between the structural stability of different tropomyosins, their endolysosomal degradation patterns and T-cell reactivity.

**Methods:** We investigated the differences between four tropomyosins - the major shrimp allergen Pen m 1 and the minor allergens Der p 10 (dust mite), Bla g 7 (cockroach) and Ani s 3 (fish parasite) - in terms of IgE binding, structural stability, endolysosomal degradation and subsequent peptide generation, and T-cell cross-reactivity in a BALB/c murine model.

**Results:** Despite their conserved primary structure and consequent IgE co-reactivity, the invertebrate tropomyosins displayed different protein stabilities. Pen m 1 and Ani s 3, but not Der p 10 and Bla g 7 elicited differential melting temperatures that were pH-dependent. Endolysosomal experiments demonstrated differential degradation, as a function of stability, generating different peptide repertoires. Pen m 1 T-cell clones, with specificity for sequences highly conserved in all four tropomyosins, did not proliferate with Der p 10, Bla g 7 and Ani s 3, indicating that these peptides were not naturally produced for other invertebrate tropomyosins.

**Conclusions:** Our data suggest that, although invertebrate tropomyosins exhibit a high degree of IgE cross-reactivity due to conserved B-cell epitopes, they do not necessarily share identical cross-reactive T-cell epitopes. This is likely due to differential endolysosomal processing as a function of different structural stabilities.

## 1. Introduction

The tropomyosin protein family is one of the largest allergen families containing over 60 identified and characterized allergens.^1^ Tropomyosin exhibits a high degree of structural conservation between species.^2^ Several studies have shown clinical cross-reactivity between crustaceans, mollusks, insects, mites and nematodes is due mainly to shared IgE (B-cell) epitopes of tropomyosin.^3–5^ However, there is a lack of understanding whether this high degree of structural and sequence conservation among tropomyosins would also lead to cross-reactive T-cell epitopes. Currently, some T-cell epitopes have only been elucidated for shrimp tropomyosin with little or no data for other allergenic sources, making analysis of T-cell cross-reactivity challenging.^6–8^

In the tropomyosin family, crustacean and mollusk shellfish tropomyosins are major allergens, particularly shrimp tropomyosin (Pen m 1), with more than 80% of shrimp-allergic patients being sensitized to this allergen. Some other invertebrate tropomyosins are considered only as minor allergens. Although allergen abundance may contribute to sensitization potential, differences in structural features or stability may also play a role.

Tropomyosins are alpha-helical coiled-coil proteins that generally exist as a dimer, and show high heat stability and resistance to pepsin digestion.^9,10^ Recent studies have demonstrated that structural and fold stability of allergenic proteins have a direct impact on their allergenicity through differential endolysosomal degradation and subsequent generation of allergen-derived peptides for MHC Class II presentation on antigen presenting cells, as shown in the case of Bet v 1, a major birch pollen allergen.^11^ Whether structural differences influence allergenicity and cross-reactivity of the various tropomyosins, remains unclear.

In this study, four allergenic tropomyosins, from shrimp (Pen m 1), house dust mite (Der p 10), cockroach (Bla g 7) and Anisakis (food borne parasite) (Ani s 3) were investigated for their structural properties and stability, as well as endolysosomal degradation and generation of peptides. Finally, T-cell cross-reactivity of these tropomyosin-derived peptides was assessed in a murine model. The tropomyosins, although heat-stable, showed different melting temperatures that were pH-dependent. Degradome analysis demonstrated that all four tropomyosins were degraded differentially leading to a non-identical repertoire of peptides. An absence of T-cell cross-reactivity of peptides generated from different tropomyosins was demonstrated in the murine model using Pen m 1 peptide-specific T-cell clones. Thus although highly conserved invertebrate tropomyosins may share IgE epitopes leading to clinical cross-reactivity, cross-reactive tropomyosin-specific T cells may not necessarily be induced due to differences in structural stability and hence dendritic cell processing and peptide presentation. Our findings have implications for understanding possible modes of sensitization as well as the design of suitable tropomyosin preparations for specific immunotherapy for shrimp and related allergies.

## 2. Materials and Methods

### 2.1 Cloning, expression and purification of tropomyosins

The amino acid sequences from the open reading frames for tropomyosin were used for Black Tiger Shrimp (*Penaeus monodon*) accession number ADM34184, House Dust Mite (*Dermatophagoides pteronyssinus*) accession number ACI32128, German Cockroach (*Blattella germanica*) accession number AAF72534.1, and Anisakis parasite (*Anisakis simplex*) accession number Q9NAS5. Detailed methods for recombinant protein expression and purification are outlined in the **Appendix S1**.

### 2.2 Patient recruitment and IgE binding analysis

To analyze IgE binding to the different tropomyosins, seventeen shellfish-allergic patients were recruited at The Alfred Hospital Allergy Clinic, Melbourne, Victoria, Australia with a positive serum shrimp-specific IgE (≥ 0.35 kU/L; ImmunoCAP®, Phadia, Uppsala, Sweden). Serum from a non-atopic donor recruited at the Translational Research Facility, James Cook University was used as a negative control (**Supplementary Table S1**). Written informed consent was obtained from all participants, and patient anonymity was preserved. Ethics approval was obtained from the Ethics Committees of James Cook University (Project numbers H4313 and H6829), The Alfred Hospital (Project number 192/07) and Monash University (MUHREC CF08/0225). IgE recognition of the different tropomyosins was investigated using grid immunoblotting^12^ as described in **Appendix S1**.

### 2.3 Biophysical characterization of allergenic tropomyosins

The alpha-helical structure and thermal denaturation of tropomyosins were compared and analyzed using circular dichroism (CD) spectroscopy. The effects of different pH conditions on protein melting temperatures was analyzed using Differential Scanning Calorimetry (DSC) and Differential Scanning Fluorimetry (DSF). Protein molecular mass under different pH conditions was determined using size exclusion chromatography coupled to multi-angle light scattering (SEC-MALS). The detailed methodology for biophysical characterization of tropomyosins is given in the **Appendix S1**.

### 2.4 Endolysosomal degradation of tropomyosins

Endolysosomal degradation assays were performed as described previously.^13^ Briefly, 5 μg of purified protein was mixed with 8 μg of isolated microsomal fraction from the JAWSII cell line in 50 mmol/L citrate buffer (pH 5.2 or pH 4.5) and 2 mmol/L dithiothreitol. Bet v 1 was used as a non-tropomyosin control allergen. Degradation was monitored over time using SDS-PAGE and Coomassie staining. The pool of peptides generated in the degradation assay was assessed by mass spectrometry using a Q-Exactive Orbitrap Mass Spectrometer (Thermo Fisher Scientific, Bremen, Germany) and nano-HPLC (Dionex Ultimate 3000, Thermo Fisher Scientific). Detailed methods are provided in **Appendix S1**. To visualize the digestion-generated peptide data, the peptide sequences were mapped against the full length amino acid sequences of the respective tropomyosins using MS tools.^14^ The speed and intensity of peptide generation was visualized using plot.ly.

### 2.5 Generation of Pen m 1 overlapping peptide library and mapping of murine T-cell epitopes of Pen m 1

To map the T-cell epitopes of Pen m 1, an overlapping peptide library was generated with 15-mer peptides with an offset of three amino acids spanning the entire length of Pen m 1 (Mimotopes, Melbourne, Australia). BALB/c mice were immunized with whole Pen m 1 and the splenocyte proliferation assay was performed using the overlapping peptide library. T-cell reactive regions were mapped, based on CFSE-based T-cell proliferation and IL-2 release as correlates. The detailed methodology is provided in the **Appendix S1**.

### 2.6 Generation of Pen m 1 conserved region-specific T-cell hybridomas

Based on T-cell proliferation data for the Pen m 1 peptide library, two peptide sequences were chosen for the generation of Pen m 1-specific T-cell hybridomas. These regions (peptides 67 ^192-^VVGNNLKSLEVSEEK^-213^ and 82 ^244-^RSVQKLQKEVDRLED^-258^) were chosen based on a) positive T-cell proliferation to these peptides and b) high amino acid sequence similarity to corresponding regions in Der p 10, Bla g 7 and Ani s 3 (Supplementary figure S1). The Pen m 1-specific hybridomas were used as a tool to investigate whether the different tropomyosins share cross-reactive T-cell epitopes at these regions. The detailed methodology for peptide immunization and generation of hybridomas is provided in the **Appendix S1**.

### 2.7 Processing and T-cell reactivity of tropomyosin homologues using Pen m 1-specific T-cell hybridomas

To investigate whether dendritic cell processing of the different tropomyosins results in the presentation of Pen m 1-crossreactive T-cell epitopes, co-cultures of Pen m 1-specific T-cell hybridoma clones with GM-CSF grown bone marrow dendritic cells (BMDCs) and serial dilutions of the four tropomyosins were set up in 96-well U-bottom plates. 10^5^ T cells per well from clones 67-1.A2 and 82-3.C5 were incubated overnight with 2×10^4^ GM-CSF BMDCs and concentrations of 50 μg/mL, 10 μg/mL, or 2 μg/mL of Pen m 1, Der p 10, Ani s 3 and Bla g 7 respectively, in triplicate. Control wells received either medium alone, peptides 67 and 82 at 10 μg/mL, or an equal volume of peptide diluent (DMSO). Culture supernatants thereof were removed and analyzed for IL-2 production as a correlate for T-cell activation using an ELISA MAX mouse IL-2 set (BioLegend, SanDiego, USA).

## 3. Results

### 3.1 Allergenic tropomyosins are highly conserved proteins with frequent IgE co-sensitization in shrimp-allergic patients

The tropomyosins selected for this study have a high degree of conservation with 70% or more amino acid sequence identity (**Figure 1A, 1B**). These allergens were expressed in a bacterial expression system and purified as recombinant proteins using affinity and size exclusion chromatography (**Figure 1C**). A multiple sequence alignment of the four tropomyosins revealed the high degree of conservation in the IgE epitopes of shrimp tropomyosin as characterized previously (**Figure 1D**).^15^ Five out of eight IgE epitopes from Pen m 1 were highly conserved in Der p 10, Ani s 3, and Bla g 7. To evaluate IgE antibody co-sensitization to the different tropomyosins, IgE grid immunoblotting was performed using sera from shrimp-allergic patients (**Figure 1E**). Thirteen out of seventeen subjects demonstrated IgE binding to all tropomyosins. Only one subject showed mono-sensitivity to Pen m 1. Interestingly, 8/17 subjects had stronger IgE binding to Der p 10 as compared to Pen m 1 based on densitometric analysis.

**Figure 1:**
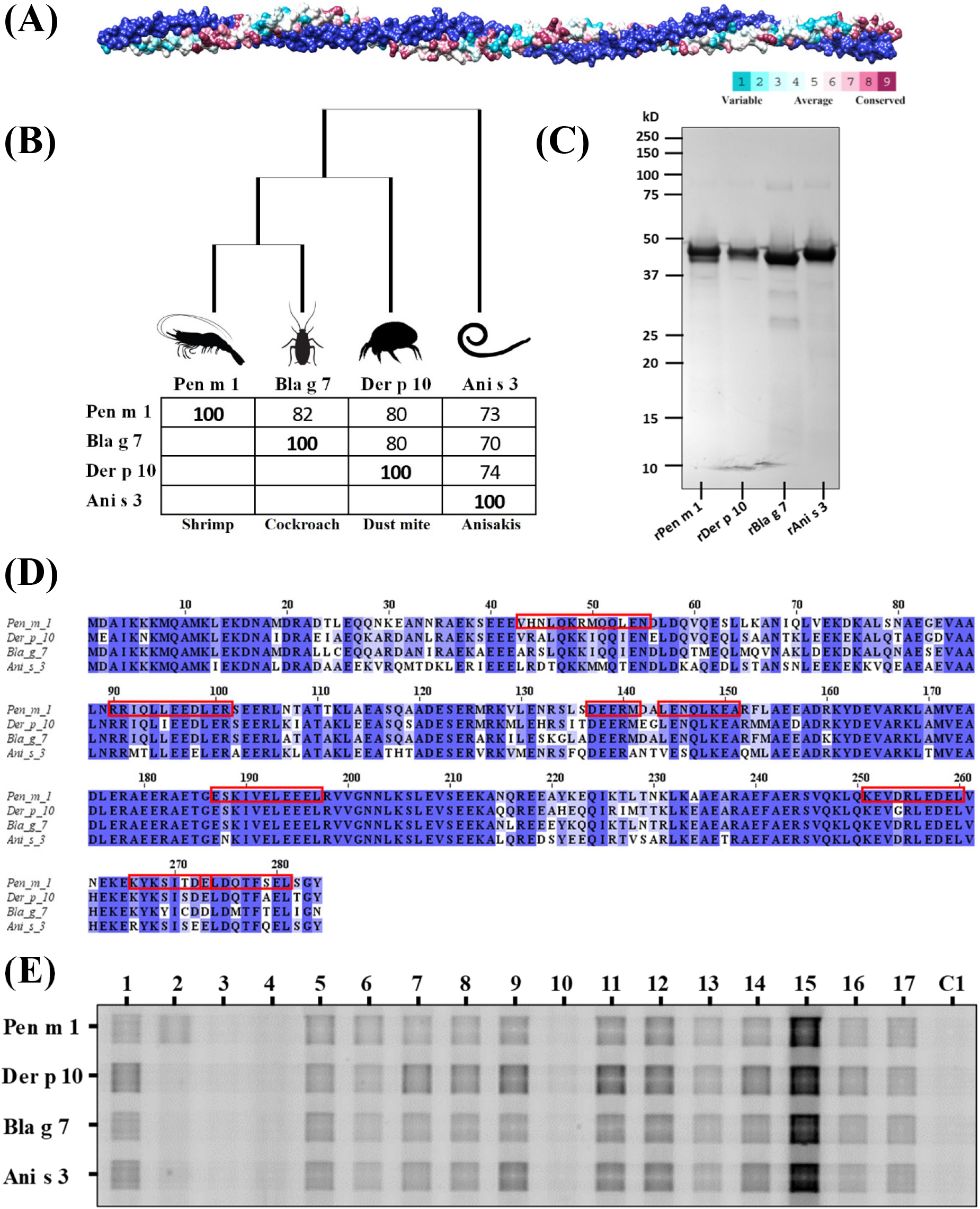
Invertebrate tropomyosins investigated in this study. (A) A homology model of tropomyosin displaying the alpha-helical coiled-coil structure and the patterns of sequence conservation. (B) A phylogenetic tree and percent identity grid for invertebrate tropomyosins investigated in this study. (C) SDS-PAGE Coomassie-stained gel profile of purified tropomyosins. (D) Multiple sequence alignment using Clustal Omega algorithm of Pen m 1, Der p 10, Bla g 7 and Ani s 3 showing conserved amino acid residues. Pen m 1 IgE binding epitopes are denoted by red boxes.^15^ (E) IgE grid immunoblotting using serum from shrimp-allergic patients (1-17) and one healthy donor (C1) to demonstrate presence or absence of IgE co-sensitization to invertebrate tropomyosins.

### 3.2 Tropomyosins have differential structural stability, and pH dependent aggregation

Using CD spectroscopy, mean residual ellipticity (MRE) at 222 nm was monitored. Pen m 1 and Der p 10 had similar melting temperatures (inflection points) of 42◻C and 44◻C respectively. Although Bla g 7 showed a higher melting temperature (63°C), loss of alpha-helical structure was initially observed from as early as 40°C (**Figure 2A, 2B**). Interestingly, Ani s 3 showed the lowest melting temperature of 33°C with nearly complete loss of alpha-helical structure by 50°C. DSF and DSC analysis of the tropomyosins confirmed the low melting temperature of Ani s 3 at neutral pH as compared to other tropomyosins (**Figure 2C, 2D**).

**Figure 2:**
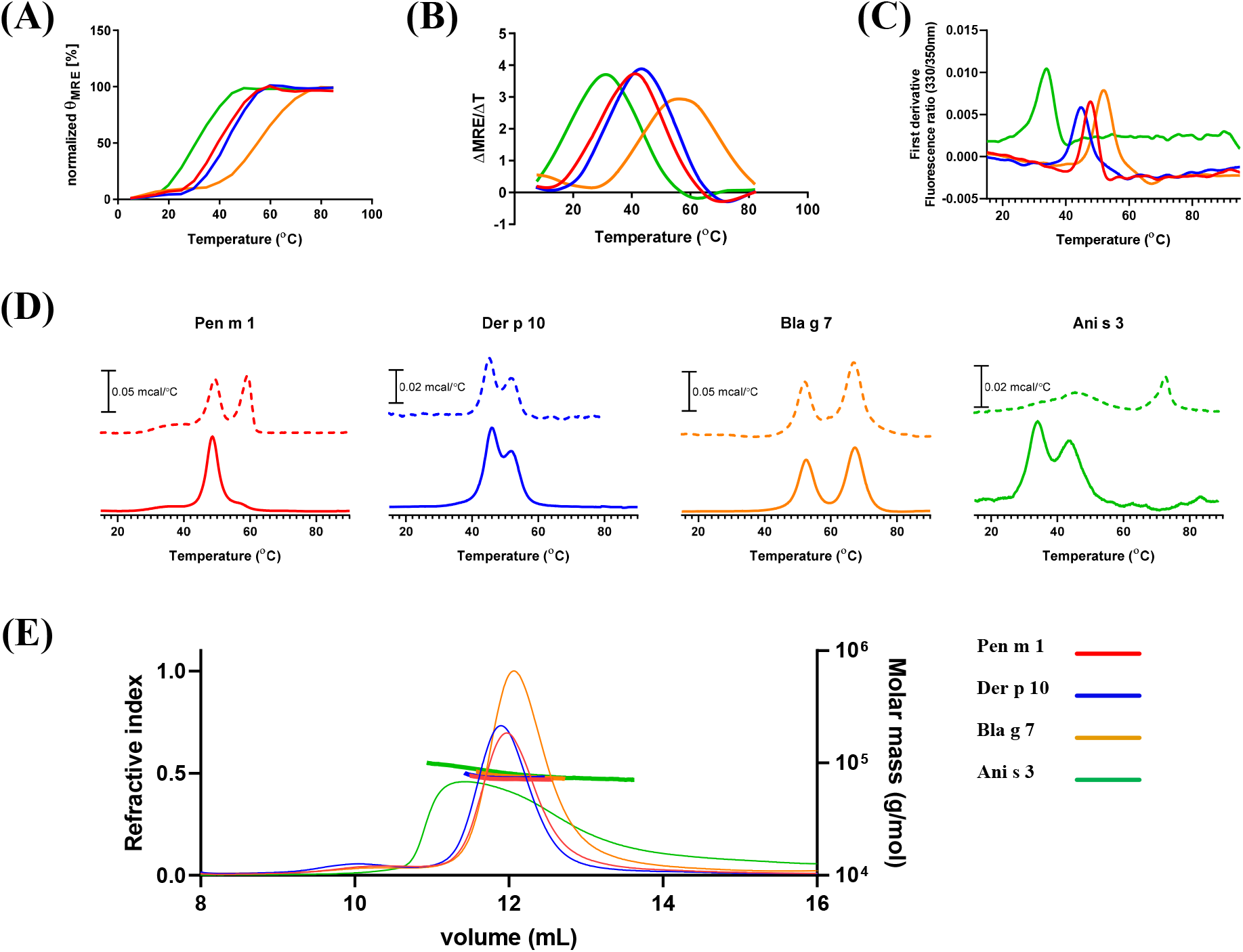
Structural characterization of tropomyosins. Analysis of thermal stability of Pen m 1, Der p 10, Bla g 7 and Ani s 3 using (A) CD spectroscopy; observing changes in MRE at 222 nm, and (B) first derivative of MRE from 15 °C to 85 °C. Analysis of thermal stability by using (C) DSF and (D) observing changes in heat capacity using (DSC) at pH 5.2 (dotted line) and pH 7.4 (solid line). (E) Analysis of tropomyosins by SEC-MALS indicated molar masses consistent with essentially full occupancy of dimer at pH 7.4. The RI chromatograms on the left axis (thin lines) while the evaluated molar masses shown with the right axis (thick horizontal lines)

The effect of two different pH conditions on melting temperatures was analyzed by DSC (**Figure 2D**). The specific heat capacity (Cp) change during protein denaturation (mcal/°C) was measured from 10°C to 85°C for the invertebrate tropomyosins at pH 5.2 and pH 7.4 in independent experiments. Pen m 1 showed a slight 2°C increase in melting temperature and an additional transition at 60°C at pH 5.2 as compared to pH 7.4. Der p 10 and Bla g 7 did not show any shifts in melting temperatures when analyzed under acidic compared with neutral conditions. Interestingly, Ani s 3 showed a large shift to 73°C at pH 5.2 from 33°C at pH 7.4.

Aggregation states of tropomyosins under acidic (pH 5.2) and neutral (pH 7.4) conditions were assessed using SEC-MALS. At neutral pH, all four tropomyosins existed as dimers as indicated by the molecular weight calculated from the peaks during size exclusion chromatography (**Figure 2E**). Ani s 3 showed a broader peak as compared to other tropomyosins. SEC-MALS was performed as a constant temperature of 25°C, and from melting temperature analysis, it was evident that Ani s 3 would have already started unfolding at this temperature, and thus we are seeing a complex equilibria of folded and unfolded protein. At pH 5.2 however, the protein peaks were absent indicating that the tropomyosins were highly charged and/or in an aggregated state, which resulted in their retention in the pre-filter or strong binding to the column matrix (data not shown).

### 3.3 Allergenic tropomyosins demonstrate differential pH-dependent protease-mediated degradation

The endolysosomal degradation assay was performed to compare the peptide profiles generated from tropomyosins in the endolysosomal compartment of antigen-presenting cells. The four tropomyosins were incubated with the microsomal fraction from JAWSII dendritic cells under different pH conditions (pH 5.2 and pH 4.5) that mimicked the increasingly acidic environment of the endosomal/lysosomal compartments (**Figure 3**).

**Figure 3:**
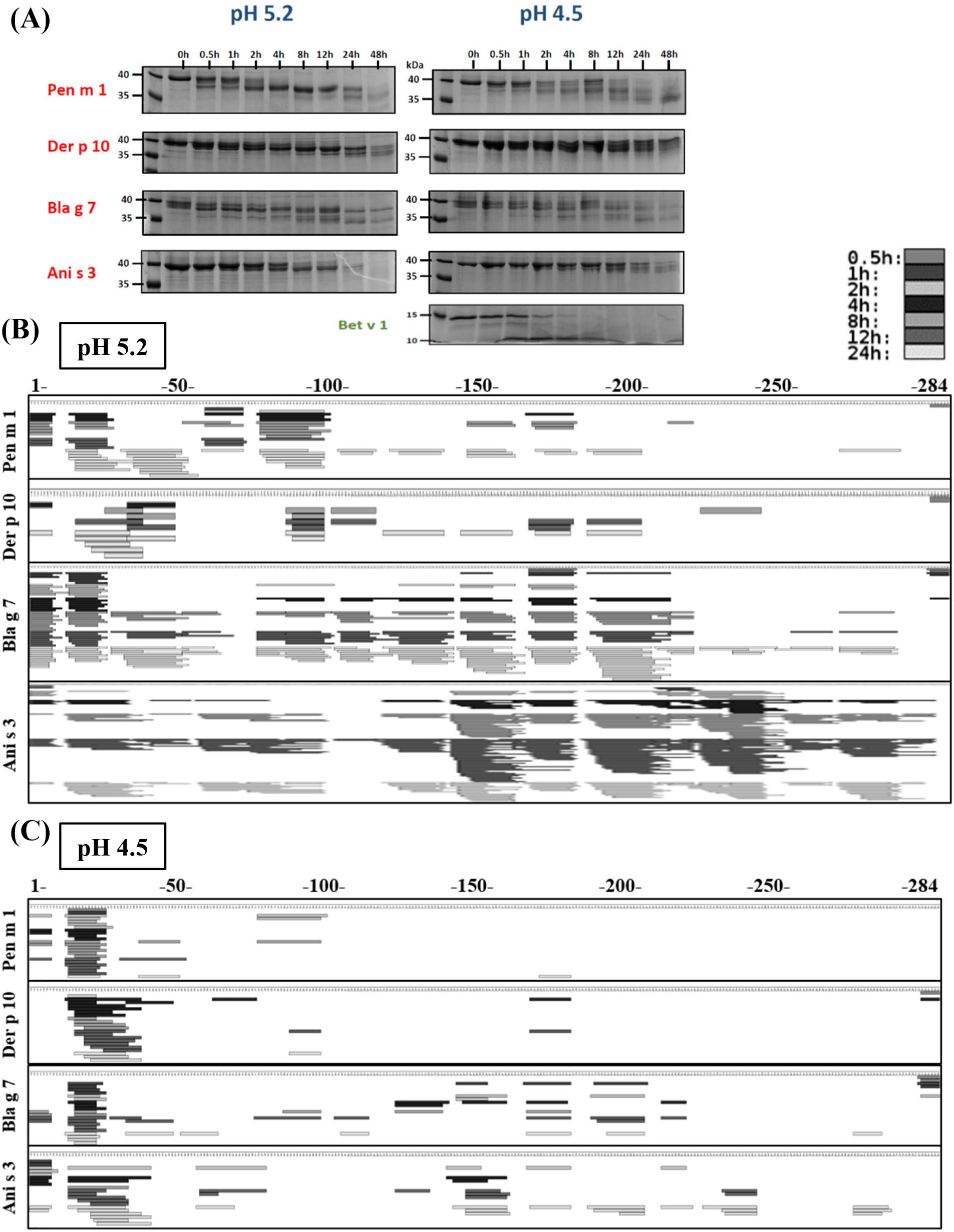
**Endolysosomal degradation of tropomyosins** using the microsomal fraction of murine dendritic cells (JAWS II) to mimic degradation in antigen presenting cells. Coomassie-stained SDS-PAGE gel of time-dependent digestion products of tropomyosins over 48 hours; lane 1 indicates molecular ladder (A). Mass spectrometric sequencing of endosomal digestion-generated tropomyosin peptides mapped against the whole amino acid sequence (numbered 1-284) for tropomyosins under acidic condition at pH 5.2 (B) and pH 4.5 (C). The time points at which peptides were generated are depicted in shades of grey (see key).

At pH 5.2, native sized Pen m 1, Der p 10 and Bla g 7 were degraded within 8 hours as observed using SDS-PAGE. However a stable 35-37 kDa fragment was visible in these cases. In contrast, Ani s 3 was completely degraded within 48 hours. At pH 4.5, degradation was slower. Der p 10 in particular was stable to enzymatic digestion at this lower pH, with a native-sized protein persisting even after 48 hours (**Figure 3A**). Interestingly, the degradation resulted in an incremental decrease in molecular weight of the tropomyosins over time at both pHs, suggesting progressive degradation from the N- and/or C-terminal ends of the coiled-coil proteins. In most cases, a residual fragment >35 kDa in size was observed. Tropomyosins in general were more resistant to endolysosomal degradation, particularly Pen m 1, as compared to the non-tropomyosin control, Bet v 1 (**Figure 3A**).

Mass spectrometry-based peptide sequencing was used to further assess the pattern of peptide generation during degradation at pH 5.2 and pH 4.5 (Figure 3B and 3C). At pH 5.2, a higher amount of digestion-derived peptides were observed than at pH 4.5 for all tropomyosins. For Bla g 7 and Ani s 3, a marked increase in peptide generation was observed after 4 hours at pH 5.2, but Pen m 1 and Der p 10 were relatively resistant to endosomal degradation even at pH 5.2 with peptides being generated at later time points (greater than 8 hours). Peptides were generated in the early time-points at the N-terminal side for all four tropomyosins as predicted using SDS-PAGE analysis, although the pattern of peptide generation was different at longer time-points particularly for Bla g 7 and Ani s 3.

The speed and intensity of peptide generation correlated with structural stability of the tropomyosins at both pH 5.2 (**Figure 4**) and pH 4.5 (**Supplementary figure 2**). Ani s 3 and Bla g 7 showed faster uncoiling of the alpha-helical structure during temperature-dependent denaturation at lower pH as shown by DSC and CD analysis, correlating with more rapid peptide generation than for the other tropomyosins.

**Figure 4:**
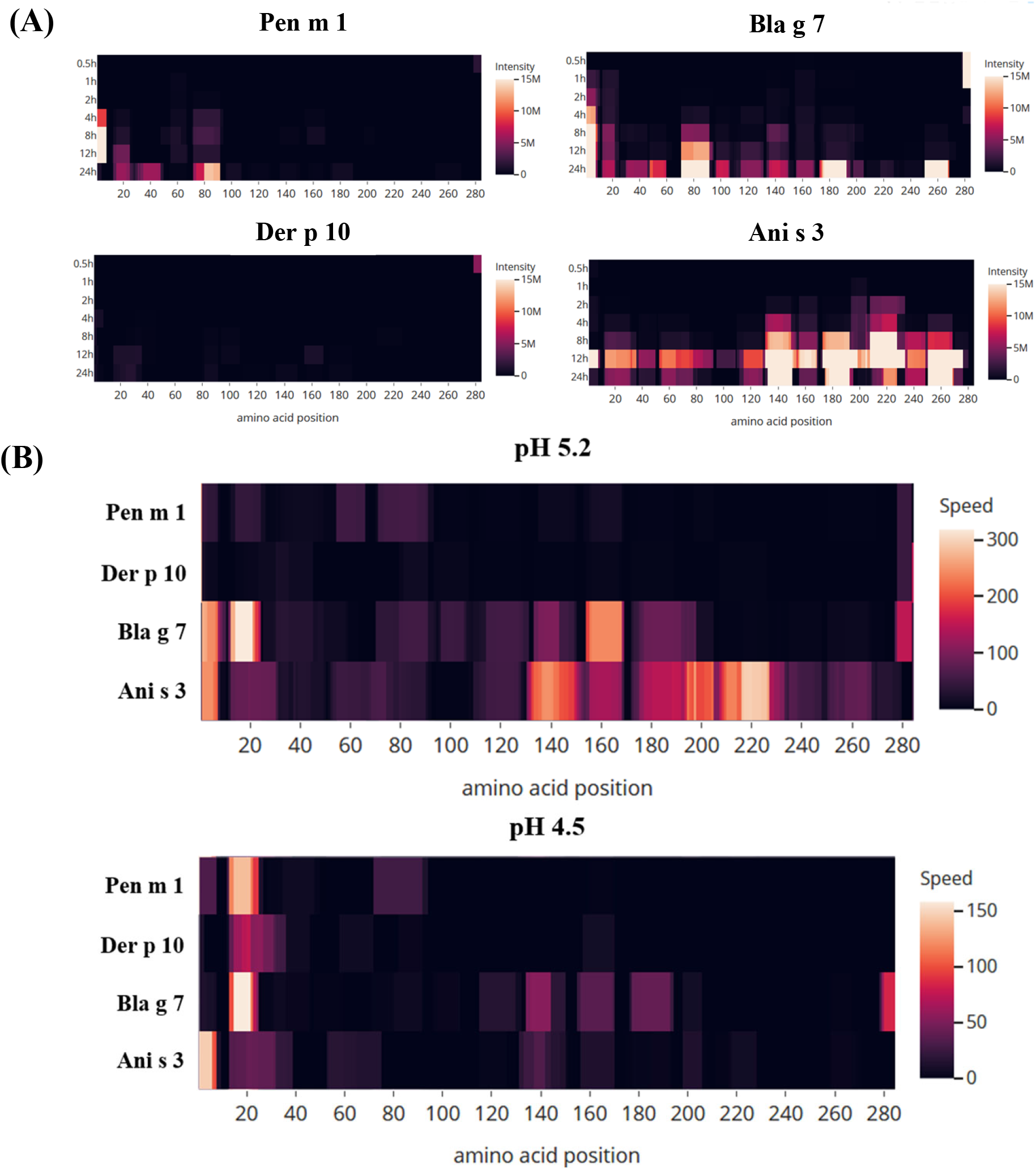
Peptide heat map depicting the intensity at pH 5.2. (A) and speed (B) of generation of tropomyosin-derived peptides during the endolysosomal degradation assay under acidic conditions. The heat map indicates the position and abundance of the generated peptides aligned against the full-length amino acid sequence of the different tropomyosins.

### 3.4 The dominant T cell epitope of Pen m 1 shows no cross-reactivity with highly homologous invertebrate tropomyosins

T-cell reactive regions of Pen m 1 were first mapped by assessing proliferation and IL-2 release (Figure 5A, 5B) from Pen m 1-immunized mouse splenocytes cultured with an overlapping Pen m 1 peptide library. T-cell reactivity was generally greater to C-terminal Pen m 1 peptides. Two T-cell reactive Pen m 1 peptides with identical or near identical sequences for corresponding regions in Der p 10, Bla g 7 and Ani s 3 (67, aa 199-213, and 82, aa 244-258) were selected for T-cell hybridoma generation (**Supplementary Figure 1**). To assess whether the invertebrate tropomyosins exhibited T-cell cross-reactivity at these conserved regions, the specific T-cell hybridomas were cultured with all four tropomyosins in separate experiments in the presence of mouse GM-CSF BMDCs as antigen presenting cells, and T-cell proliferation was assessed by IL-2 release (**Figure 6**).

**Figure 5:**
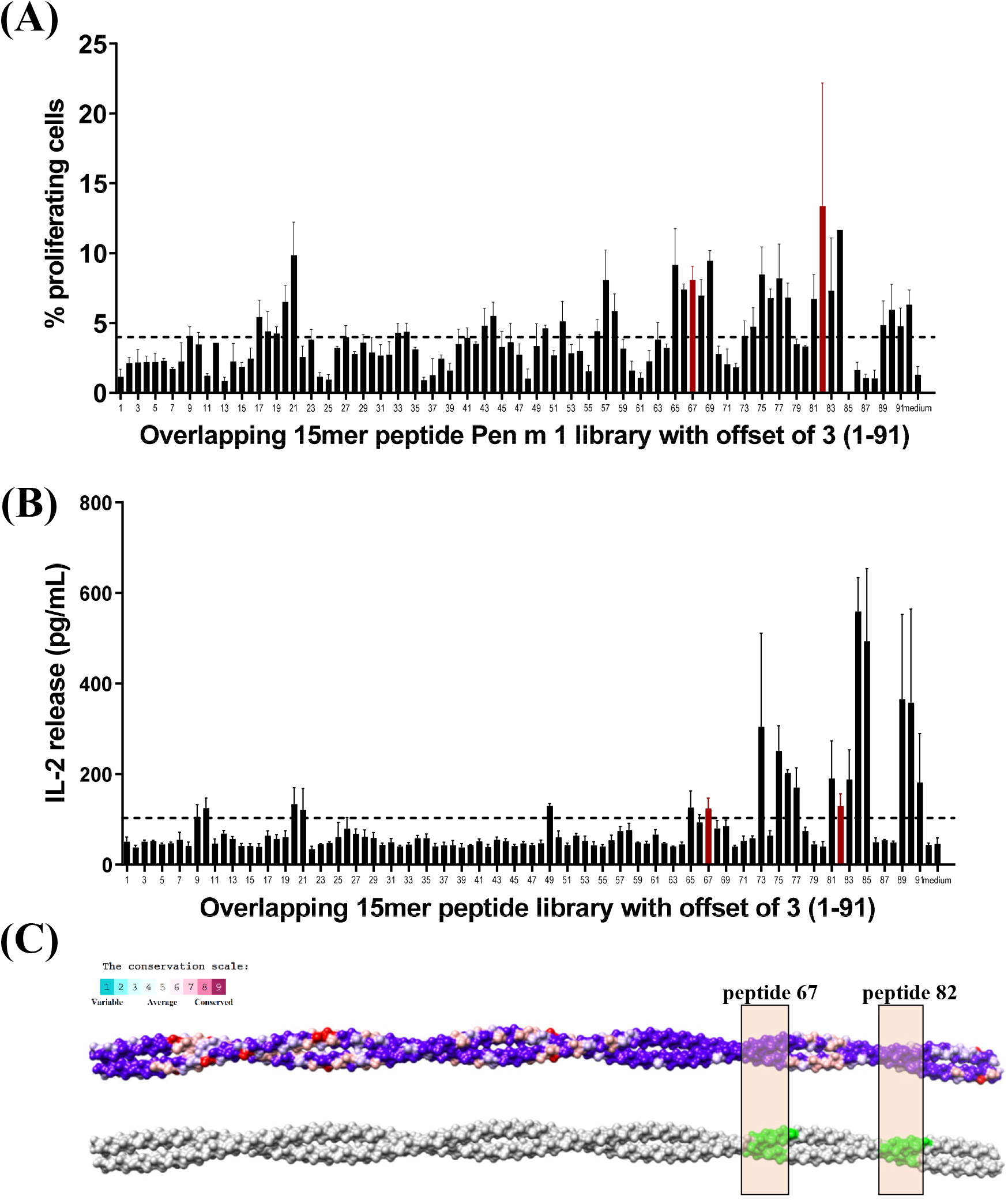
Murine T-cell epitope mapping of Pen m 1. Immunoreactive regions of Pen m 1 were mapped using overlapping 15-mer peptides with an offset of 3 amino acids. (A) Proliferating T-cells were analyzed using CFSE dye-dilution method. B) IL-2 release in the supernatant was measured using ELISA. Data are shown as mean with standard error of mean (SEM) for three replicate cultures. The cut-off of three standard deviations above mean reactivity of medium-only control wells is indicated by the dotted line (A, 3.98% proliferating cells or B, 103.2 pg/mL IL-2). (C) A model of tropomyosin representing the amino acid conservation between Pen m 1, Der p 10, Bla g 7, and Ani s 3 generated using Consurf conservation model in Chimera; the purple regions indicate highly conserved regions. The green shade indicates the two T-cell reactive regions of Pen m 1 selected for this study that are also highly conserved among the four allergenic tropomyosins

**Figure 6:**
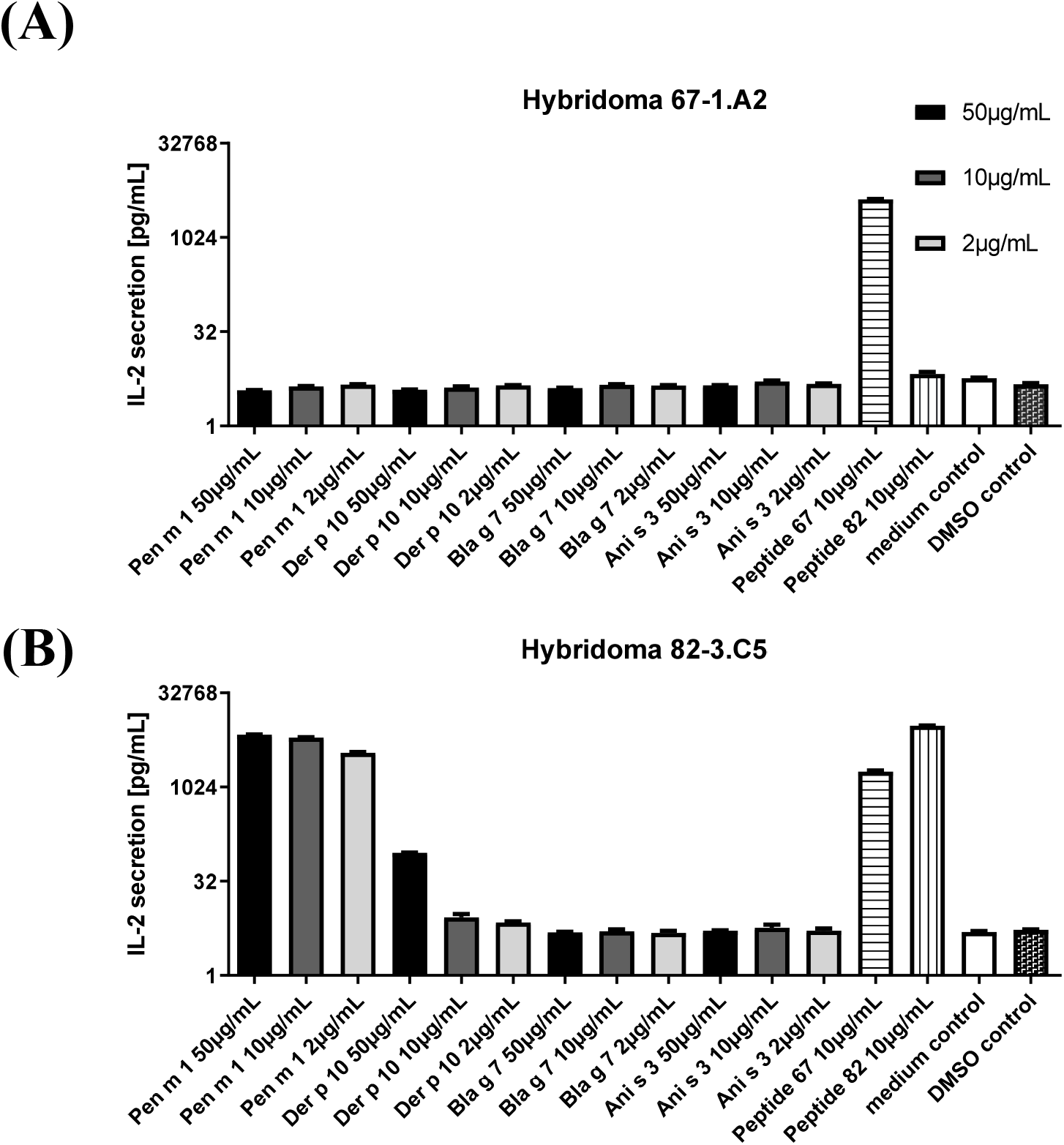
Murine T-cell cross reactivity to allergenic tropomyosins. T-cell cross-reactivity of Pen m 1, Der p 10, Bla g 7, and Ani s 3 to T cell hybridoma clones specific for peptide 67 and 82 was analyzed by measurement of IL-2 production upon exposure to protein, peptide controls or medium. Data are shown as mean with standard error of mean (SEM) for three replicate cultures.

Although peptide 67 (positive control) induced strong proliferation of clone 67-1.A2, none of the four tropomyosins or peptide 82 did. The clone 82-3.C5 (generated against peptide 82) showed reactivity to both peptide 67 and peptide 82. This clone showed a proliferative response to Pen m 1 at all three tested concentrations, while Der p 10 elicited a weak response only at the highest concentration tested and Bla g 7 and Ani s 3 did not induce any T-cell proliferation. Notably, these regions are highly similar (peptide 67: 100% for all three homologues; peptide 82: 93% for Der p 10, 100% for Bla g 7 and Ani s3) among all four tested tropomyosins, and yet cross-reactivity was not observed.

## Discussion

Shellfish allergy, particularly to shrimps, affects more than 3% of the population and is frequently associated with life-long sensitivity and severe allergic reactions on exposure.^16–18^ Tropomyosin, a major allergen, has been the central focus of immunodiagnostic and therapeutic developments for shrimp allergy. Tropomyosins are known to be highly conserved structural proteins, which forms the molecular basis for their IgE antibody cross-reactivity. Tropomyosins have been key players in our understanding of clinical cross-reactivity in shrimp-allergic patients on exposure to other invertebrate sources such as insects ^19,20^, mites ^21^, nematodes ^22^ and more recently, even vertebrates such as fish.^23,24^ Tropomyosin is the primary sensitizer in various crustacean species that leads to a strong Th2 response in humans and in small animal models ^25^. However, it remains to be elucidated whether exposure to other invertebrate tropomyosins acts as a potent activator of shrimp-specific T cells and if shrimp tropomyosin (Pen m 1) shares identical cross-reactive T-cell epitopes with other allergenic tropomyosins due to its highly conserved sequence and heat-stable structure.

From a clinical perspective, although avoidance of any shrimp-based foods can prevent further sensitization and IgE-mediated reactions in shrimp-allergic patients, it is not known whether environmental exposure to dust mite or insect derived tropomyosin via inhalation, or to the fish parasitic nematode Anisakis tropomyosin via ingestion, could lead to further priming and re-stimulation of a shrimp-specific allergic response. T-cell cross reactivity has recently been demonstrated for nut allergy, where cashew-specific CD4+ T cells from cashew-allergic subjects can cross-react with tree nut allergens such as hazelnut and pistachios due to homologous T-cell epitopes with high amino acid identity.^26^ Similarly, pollen-related food allergens were shown to induce Bet v 1-specific T-cell responses in pollen-allergic subjects, which might perennially boost IgE levels out of the pollen season due to consumption of such foods.^27^

In this study, we sought to understand the fundamental relationship between allergenic invertebrate tropomyosins in terms of their structural stabilities, endosomal degradation patterns and T-cell reactivity. Using an animal model, we investigated whether Pen m 1-specific T cells could cross-react with other invertebrate tropomyosins due to conserved T-cell epitope regions.

Four tropomyosins were investigated in this study: Pen m 1 (shrimp), Der p 10 (house dust mite), Bla g 7 (cockroach) and Ani s 3 (Anisakis), all of which have been meticulously investigated for their allergenicity and registered on the IUIS allergen database.^28^ Through our detailed structural analysis, we demonstrate here that these tropomyosins, despite their conserved primary structure, exhibit different stabilities as reflected in their melting temperatures, while all are known to withstand heat-processing and retain IgE binding capacity. Ani s 3 showed a remarkably low melting temperature, which may be a physiological property of a fish parasite protein that is adapted to a cooler environment. In contrast, an increase in melting temperature was observed for Pen m 1 and Ani s 3 under acidic pH conditions. Importantly both Pen m 1 and Ani s 3 are food-derived allergens giving them added stability under the acidic conditions of the stomach. Similarly, allergens are subjected to increasingly acidic conditions after uptake by antigen presenting cells in the early to late endolysosomal compartments.^11,29^ This variation in stability at acidic pH indicates that the different tropomyosins might be degraded differentially during antigen presentation.

During allergic sensitization, allergens are taken up by antigen presenting cells (APCs) and degraded inside the endolysosomal compartment by proteases including cathepsins C, H, Z and AEP.^13^ Specific peptides that are generated during this degradation process are loaded onto MHC class II molecules and displayed on the cell surface for presentation to CD4+ T cells, eventually resulting in a Th2 cytokine response and subsequent generation of allergen-specific IgE. To establish whether the different invertebrate tropomyosins are degraded and processed differently during this stage, the endolysosomal degradation assay was performed at pH 5.2 and pH 4.5 to mimic the early and late endosomal compartments of APCs. For the first time, we demonstrate that tropomyosins are degraded differentially and at different speeds depending on pH conditions. The speed and intensity of peptide generation was higher for Ani s 3 and Bla g 7 as compared to Pen m 1 and Der p 10. Ani s 3 in particular showed a low melting temperature, which may have led to enhanced unwinding or fraying of the coiled-coil structure, as corroborated by DSC and SEC-MALS, giving way for more efficient degradation of the peptide bonds. We showed that the generated peptide sequences, as well as the rate and intensity of peptide generation, differed between tropomyosins as a function of their pH-dependent structural stability.

To further investigate whether this differential degradation of tropomyosins would translate to differential T-cell reactivity *in vivo*, murine T-cell clones were generated from mice immunized with Pen m 1 peptide 67 (aa 199-213) and peptide 82 (aa 244-258). These regions were T-cell reactive in Pen m 1, and highly conserved among all four tropomyosins. However when all four tropomyosins were cultured with these T-cell clones, Bla g 7 and Ani s 3 did not induce any significant proliferation. Der p 10 induced some IL2 release but only at the highest tested concentration. Clone 67-1.A2 did not exhibit any proliferation even on exposure to whole Pen m 1. Surprisingly, peptide 67 was able to induce proliferation in T-cell clone 82.3.C5 indicating T-cell receptor (TCR) cross-reactivity between these two internal regions of Pen m 1. A single TCR has been shown to recognize more than one specific peptide.^30,31^ We conclude that peptide 67 was only identified in the initial screening using whole Pen m 1 due to its crossreactivity with the immunodominant region 244-254, as T cell clones specifically raised against peptide 67 fail to be stimulated by all tested tropomyosins. Unlike tree-nut allergen^26^ or pollen allergen^32^ T-cell cross-reactivity, the tropomyosin family of allergens may not share identical cross-reactive T-cell epitopes, and this may be attributed to their different structural stabilities. However, the alpha-helical coiled-coil structure imparts features unique to tropomyosins, namely a heptad repeat amino acid sequence.^33,34^ This feature makes it possible to have regions which have different amino acid sequences but share similar classes that would act as anchor points to the peptide groove of MHC class II molecules. This could result in non-homologous “non-identical” cross-reactive T-cell epitopes ‘within’ and ‘among’ tropomyosins. This warrants further investigation.

A point to note in this study is that inhaled tropomyosins are exposed to a different environment than those taken up via the gastrointestinal tract. Inhaled food allergens, or highly similar inhalant allergens, may induce allergic sensitization and eventually lead to ingestion-induced food allergy. The varying protein stability and different biophysical behavior of tropomyosin that we demonstrate in this study, may play a crucial role in the subsequent IgE sensitization via these two different routes as seen in the case of these recently termed ‘class 3’ allergens among shellfish processing workers.^35^

In summary, invertebrate tropomyosins do not share similar stabilities, despite a high degree of primary sequence conservation. Stability difference may be the prime reason for differential degradation and antigen presentation of various allergenic tropomyosins, leading to generation of non-identical T-cell epitopes. In our current understanding, although a high degree of IgE cross-reactivity and co-sensitization is observed among different allergenic tropomyosins, they may not be able to co-stimulate a T-cell response. However, further investigation including allergic subjects, and a wider range of T-cell epitopes is required to fully evaluate and understand T-cell cross-reactivity among various allergenic tropomyosins. Further understanding of the presence and dynamics of independent and cross-reactive T-cell epitopes for the tropomyosin allergen family is required for elucidating modes of sensitization, cross-sensitization as well as development of effective T-cell targeted immunotherapy for shrimp and related allergies.

## Acknowledgements

This project was funded by the Austrian Science Fund (FWF) project P26997_B13, Allergy and Immunology Foundation of Australasia (AIFA), and National Health and Medical Research Council (NHMRC) project grant GNT1086656. SK is an NHMRC Peter Doherty Early Career Research Fellow (GNT1124143).

---

**Supplementary Figure S1: A schematic diagram representing the 91 overlapping Pen m 1 peptides**. Peptides that showed positive T-cell proliferation (red), IL-2 release (yellow) or both (orange) are shown. Multiple sequence alignment of Pen m 1, Der p 10, Bla g 7 and Ani s 3 is shown with non-identical amino acids highlighted in green. The amino acid sequences of peptides 67 and 82, that were chosen for generating the T-cell hybridomas, are highlighted in blue.

**Supplementary Figure S2: Peptide heat map depicting the intensity of tropomyosin-derived peptides** generated at different times during the endolysosomal degradation assay under acidic conditions at pH 4.5. The heat map indicates the position of the generated peptides against the aligned full length amino acid sequences of the different tropomyosins

