## Supplementary material for "Effect of structural stability on endolysosomal degradation and T-cell reactivity of major shrimp allergen tropomyosin": Table S1

**Supplementary Table S1:** Details on serum-based shrimp-specific and HDM-specific IgE levels and associated clinical symptoms for shellfish-allergic patients recruited for this study

| **Slot** | **IgE**  **(kU/L)** | **shrimp RAST (kUA/L)** | **HDM RAST (kUA/L) or SPT** | **Clinical symptoms** |
| --- | --- | --- | --- | --- |
| 1 | 199 | 3(10.10) | 8 mm (SPT) | Anaphylactic, eyes swelling, oral itch |
| 2 | 3401 | 3(6.65) | 6(>100) | Urticaria, Angioedema |
| 3 | 127 | 2(1.65) | 0(0.09) | Lip swelling on contact |
| 4 | 242 | 2(1.32) | 11 mm (SPT) | Urticaria on handling |
| 5 | 288 | 3(9.82) | 2(2.66) | Facial oedema, red rash |
| 6 | 183 | 3(6.84) | 4(31.70) | NR |
| 7 | 579 | 3(3.63) | 3(5.03) | Oral/ hand contact |
| 8 | 266 | 3(17.2) | NT | Oral |
| 9 | 227 | 3(9.81) | 3(16.80) | Angiodema |
| 10 | 1946 | 3(9.50) | 3(14.10) | Oral |
| 11 | 323 | 4(21.60) | NT | Urticaria |
| 12 | 391 | 3(14.10) | 5(71.40) | Oral |
| 13 | 449 | 3(5.42) | 4(40.20) | Anaphylaxis, angioedema |
| 14 | 322 | 3(6.73) | 3(6.90) | Anaphylaxis, urticaria |
| 15 | 3424 | 6(>100) | 4(22.00) | Angiodema, hives |
| 16 | 1665 | 3(12.6) | NT | Oral itch, exercise |
| 17 | 32 | 3(5.05) | NT | Itchy and tight throat |

NT – Not tested, NR – Not recorded
