## Supplementary figures and images for "Effect of structural stability on endolysosomal degradation and T-cell reactivity of major shrimp allergen tropomyosin"

### Figure S1

**Supplementary figure S1:**

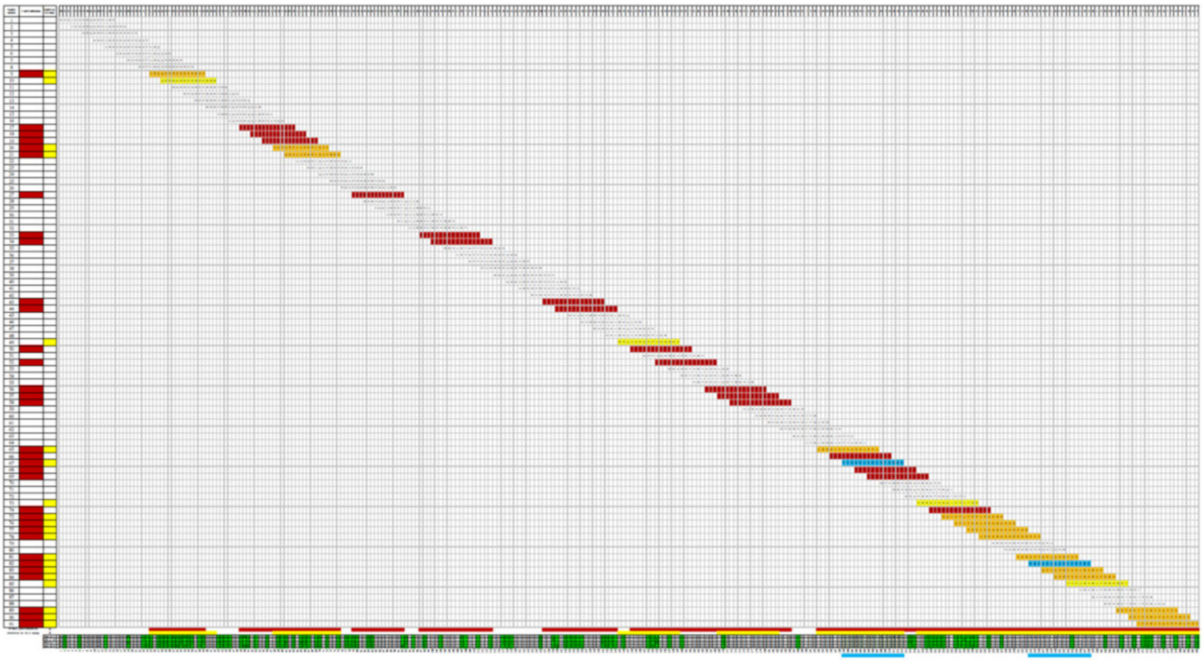

### Figure S2

#### Supplementary figure S2:

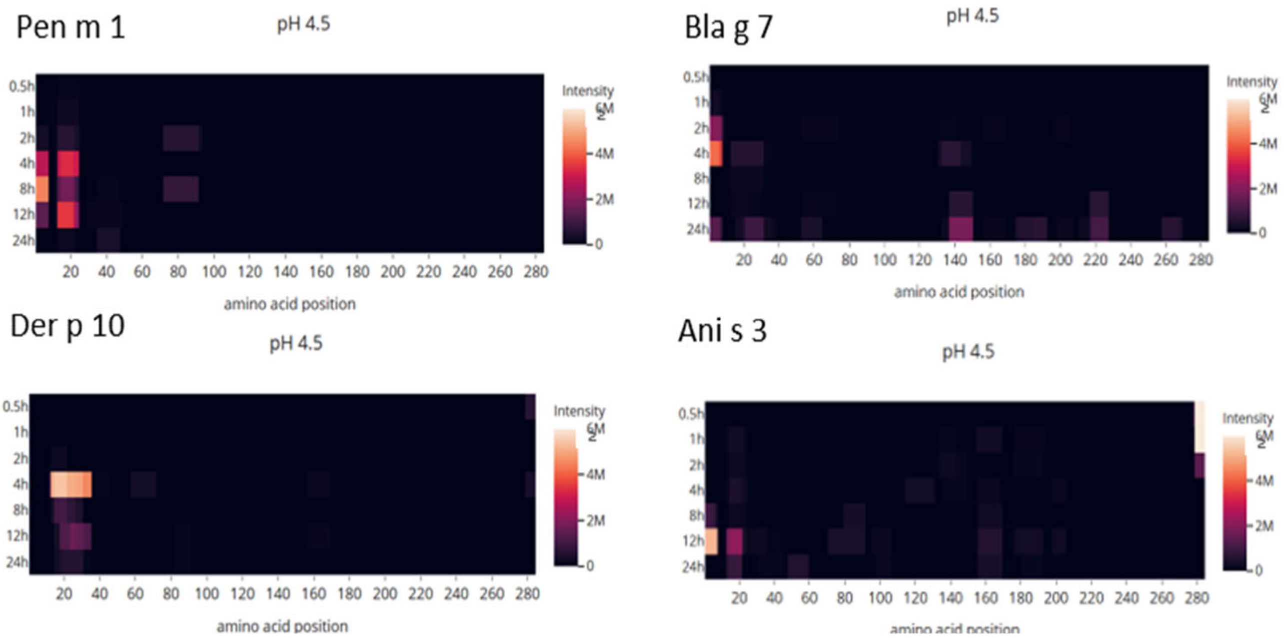
