## Appendix S1 for "Effect of structural stability on endolysosomal degradation and T-cell reactivity of major shrimp allergen tropomyosin"

**APPENDIX S1 (Supporting information)**

**Supplementary Materials and Methods**

**Cloning, expression and purification of tropomyosins**

The codon optimized DNA sequences of the open reading frames were synthesized (Life Technologies, Carlsbad, USA) and cloned into the expression vector pRSET-A (Life Technologies, Carlsbad, USA). The tropomyosins were expressed as recombinant proteins using a bacterial expression system as described previously.^1^ Briefly, chemically-competent BL21 (DE3) RIPL *E.coli* cells were transformed using these plasmids at 40ᴼC for 45 seconds and grown overnight on LB agar medium. Protein expression was induced using 0.6 mM IPTG for 4 hours at 37ͦC, and after centrifugation of the culture, the cells were washed and lysed using ultrasonication (Branson, Danbury, USA).

The recombinant proteins were purified from the bacterial lysate using IMAC affinity chromatography and then further purified by size exclusion chromatography using a Superdex 75 hiLoad column (GE Healthcare, Chicago, USA) with PBS, pH 7.2 as a mobile phase. The purified proteins were finally desalted into 50 mM ammonium bicarbonate buffer using a HiTrap desalting column (GE Healthcare, Chicago, USA), lyophilized and stored at 4ᴼC for further use.

**IgE grid-immunoblotting**

IgE recognition of the different tropomyosins was investigated using grid immunoblotting.^2^ One microgram of the purified protein in PBS, pH 7.2 was coated onto a nitrocellulose membrane for one hour. After blocking the membrane with 2% casein blocking buffer (Sigma Alrich, St Louis, USA), the membrane was incubated overnight at 4ᴼC, with shellfish-allergic patient or non-atopic control donor serum diluted 1:20 in casein blocking buffer using the surf-blotting apparatus (Idea Scientific, Minneapolis, USA). After washing with PBS/0.05% Tween-20, the membrane was incubated with rabbit anti-human IgE (Dako, Santa Clara, USA) and subsequently with goat anti-rabbit IgG conjugated to Dylight 800 (Thermo Fischer, Waltham, USA). IgE binding to tropomyosin was visualized using an Odyssey CLx infra-red imaging system (Licor, Lincoln, USA).

**Biophysical characterization of allergenic tropomyosins**

**Circular dichroism spectroscopy**

The alpha-helical structure of the tropomyosins was confirmed using circular dichroism spectroscopy. The proteins were diluted from lyophilized stocks to 0.1 mg/ml in 10 mM phosphate buffer, pH 7.2 and analyzed using a Jasco-J815 spectrophotometer (Jasco, Easton, USA) in a cell with 1 mm path length and constant nitrogen flushing. The spectra were recorded at 25ᴼC from 180 nm to 260 nm with a step resolution of 1nm at a scan speed of 50nm/min. A total of five scans were accumulated for each spectrum.

To assess the thermal denaturation of the tropomyosins, protein samples were prepared in 10 mM phosphate buffer, pH 7.4 at a final concentration of 0.1 mg/mL. A Jasco J-1500 spectrophotometer was used with a sample cell of 1 mm pathlength under constant nitrogen flushing. The changes in alpha-helical content were monitored at 222nm from 5ᴼC to 95ᴼC with 5ᴼ step-wise increments.

Tropomyosin samples were prepared from lyophilized material by dissolving directly in 50mM sodium phosphate, 150 mM NaCl pH 7.4 buffer at 1 mg/ml. Samples at pH 5.2 were first dissolved at 5 mg/ml in ultrapure water and then diluted to 1 mg/ml in a final working buffer of 50mM sodium acetate, 150 mM NaCl pH 5.2. In both cases samples were vortexed to dissolve and centrifuged at 15000 rpm for 5 minutes using a bench top centrifuge before use.

**Differential Scanning Calorimetry**

DSC of tropomyosin samples in PBS was performed using a Microcal CAP-DSC instrument (GE Healthcare) in mid feedback mode. Protein concentration was 1 mg/ml and a scanning rate of 2 °C/min between 5-105 °C was used. The repeatability of thermal denaturation was assessed by cooling and rescanning samples in the DSC. There was no evidence for reversibly refolded material producing a rescan transition indicating the thermal denaturation was an irreversible process.

**Differential Scanning Fluorimetry (DSF)**

Thermal denaturation was also followed using intrinsic protein fluorescence on a Nanotemper Prometheus instrument. Tropomyosin samples at 1 mg/ml were loaded into standard capillaries and scanned at 2 °C/min. The first derivative of the fluorescence emission ratio 350/330 nm was used to follow denaturation and the presence of aggregation was evaluated using back-reflected absorbance that is increased as any particles formed scatter light.

**Size exclusion chromatography – Multiple angle light scattering**

Tropomyosin samples at 1 mg/ml were first resolved on a Superdex-200 HR10/300 analytical gel filtration column (GE Healthcare) at 0.5 ml/min in buffer before light scattering detection on a Wyatt Heleos II 18 angle light scattering instrument coupled to a Wyatt Optilab rEX online refractive index detector (Wyatt Technology Corporation) in a standard SEC-MALS format. Protein concentration was determined from the excess differential refractive index based on 0.186 RI increment for 1 g/ml protein solution. Concentrations and observed scattered intensities at each point in the chromatograms were used to calculate the absolute molecular mass from the intercept of the Debye plot, using Zimm formalism as implemented in Wyatt’s ASTRA software.

**Endolysosomal degradation and Mass spectrometric analysis**

The pool of peptides generated in the degradation assay was assessed by SDS-PAGE/coomassie staining and mass spectrometry using a Q-Exactive Orbitrap Mass Spectrometer (Thermo Fisher Scientific, Bremen, Germany) and nano-HPLC (Dionex Ultimate 3000, Thermo Fisher Scientific). Prior to chromatography, samples were desalted with C18 ZipTips (EMD Millipore, USA) according to the manufacturer’s instructions. Chromatography was performed with an Acclaim PepMap RSLC column (75 μm x 215 cm, Dionex) using an acetonitrile gradient (Solvent A: 0.1% (v/v) formic acid; solvent B: 0.1% (v/v) formic acid/80% (v/v) acetonitrile; 5–45% B in 60 min) at a flow rate of 300 nL/min at 55°C. The HPLC was directly coupled via nano electrospray to the mass spectrometer. Capillary voltage was 2 kV, and the instrument was tuned to maximum sensitivity. For peptide assignments, a top 12 method was used with the normalized fragmentation energy at 27%. Peptides were identified using PEAKS Studio 8.5 (Bioinformatics Solutions, Waterloo, ON, Canada). Searches were conducted with single allergen sequences and only peptide hits with high confidence scores (−10lgP ≥ 35) were considered.

**Mapping murine T-cell epitopes of Pen m 1**

Female BALB/c mice (Janvier Labs, Le Genest-Saint-Isle, France) were immunized 3 times with a 2-week and a 4-week interval in between, respectively, by subcutaneous injection of 10 µg purified recombinant Pen m 1 adjuvanted with Al(OH)3 (Alu-Gel-S, Serva, Heidelberg, Germany, 50%v/v) in endotoxin-free PBS (200 µL total volume). Three weeks after the 3rd immunization, mice were killed and spleens and axillary and inguinal lymph nodes were harvested. All animal experiments were performed in accordance with EU Directive 2010/63/EU and approved by the Austrian Ministry of Education, Science and Research, permit number BMWF-66-012/0024-II/3b/2016. Following preparation of single cell suspensions, red blood cells were lysed and lymphocytes were stained with CFSE (eBioscience, San Diego, USA) according to the manufacturer’s instructions. Briefly, 5x10^7^ pooled spleen and lymph node cells from each mouse were incubated with CFSE in 10 mL warm DPBS at a concentration of 1 µM for 10 min in a 37°C water bath. Labeling was stopped by addition of 10% FCS. After two washing steps with DPBS, lymphocytes were cultured in MEM supplemented with 1 mM L-glutamine, 1% FCS, 1xPenStrep, 1 mM sodium pyruvate, 20 mM HEPES, 1x non-essential amino acids, 50 µM β-mercaptoethanol at 2x10^5^ cells per well in U-bottom tissue culture plates (Greiner, Kremsmünster, Austria) with individual peptides from the 15mer library (Mimotopes, Melbourne, Australia) with an offset of 3 amino acids (final concentration 10 µg/mL) or recombinant full-length Pen m 1 (20 µg/mL) in triplicate cultures.

After 7 days of incubation, culture supernatants (50 µL/well) were carefully removed and stored at -20°C for analysis of IL-2 production. Cells were resuspended in the remaining medium, transferred to V-bottom plates and stained in DPBS for 10 min at RT with fixable viability dye eFluor 780 (1:5000, eBioscience) followed by anti-mouse CD4 PerCP/Cy5.5 (1:200, BioLegend, SanDiego, USA, clone GK1.5) in FACS buffer (PBS, 1%BSA, 2 mM EDTA), and finally analyzed on a FACSCanto II flow cytometer (BD Biosciences, San Jose, USA). Proliferating cells among live CD4+ T cells were identified by their reduced fluorescence on the FITC channel. Measurement of IL-2 in culture supernatants as a correlate of T-cell activation was performed using an ELISA MAX mouse IL-2 set (BioLegend) according to the manufacturer’s instructions. Peptides were considered immunoreactive if the percent T-cell proliferation or IL-2 release was greater than three standard deviations above the mean of medium-only control wells.

**Peptide immunization and generation of Pen m 1 peptide-specific T-cell hybridomas**

To generate specific T-cell responses against regions with high (93-100%) amino acid conservation between Pen m 1, Der p 10, Bla g 7, and Ani s 3, which also induced T-cell reactivity, immunization with individual peptides 67 and 82 was performed. Mice were first immunized with peptide in complete Freund’s adjuvant. Emulsions were prepared in 1 mL syringes by mixing 125 µL complete Freund’s adjuvant with 125 µL PBS containing 200 µg of peptide and vigorously shaking for 6 h on a shaker. Animals received two subcutaneous injections on their backs and emulsions were distributed by massage. Immune responses were boosted twice at 10-day-intervals with peptides adjuvanted with incomplete Freund’s adjuvant. To this end, a mixture of 850 µL Bayol F and 150 µL Aracel A was prepared. 125 µL thereof were filled into the syringes followed by 125 µL PBS and 200 µg peptide. Again, the syringes were shaken vigorously for 6 h prior to subcutaneous injection. Five days after the third immunization, mice were killed and spleens, axillary and inguinal lymph nodes were harvested. Following preparation of pooled single cell suspensions for each mouse and lysis of red blood cells, lymphocytes were cultured in Opti-MEM (Gibco), 5% FCS, 1x PenStrep, with the respective peptide used for immunization at a final concentration of 3 µg/mL. One week later, peptides were removed by washing and cells incubated with 20 U/ml recombinant human IL-2 for another week. After removal of IL-2 by washing, T cells were co-cultured with syngeneic mouse GM-CSF bone marrow derived dendritic cells (BMDCs) (10:1 ratio) in the presence of 5 µg/mL of the respective peptide. On day 4, T cells were harvested and fused with mouse thymus lymphoma cell line BW5147.G.1.4 (ATCC) according to standard protocols for generation of T-cell hybridomas.^3^ The hybridomas were cloned at least three times using single cell sorting on the FACS Aria III for achieve 100% clonality.
